# Messenger RNA levels of enzymes involved in glycerolipid synthesis in the brain of the mouse and its alterations in *Agpat2*^-/-^ and db/db mice

**DOI:** 10.1101/2020.04.02.021808

**Authors:** Lila González-Hódar, Anil K. Agarwal, Víctor Cortés

## Abstract

**Aims:** Expression of genes encoding enzymes involved in glycerolipid and monoacylglycerol pathways in specific brain regions is poorly known and its impact in insulin resistance (IR) and type 2 diabetes (T2D) in the brain remains unreported. We determined mRNA levels of enzymes involved in glycerolipid synthesis in different regions of the mouse brain and evaluated their changes in two models of IR and T2D, the *Agpat2*^*-/-*^ and *Lepr*^db/db^ mice.

**Methods:** Cerebral cortex, hypothalamus, hippocampus and cerebellum were dissected from adult *Agpat2*^-/-^ mice, *Lepr*^db/db^ mice and their respective wild type littermates. Total RNA was isolated and mRNA abundance was measured by RT-qPCR.

**Key findings:** GPAT1, AGPAT1-4, LIPIN1/2, DGAT1/2 and MOGAT1 mRNAs were detected in all studied brain regions, whereas GPAT2, LIPIN3 and MOGAT2 were undetectable. Abundance of AGPATs, LIPIN1 and DGAT1, was higher in cerebellum and hypothalamus. LIPIN2 and MOGAT1 levels were higher in hypothalamus, and DGAT2 was higher in cortex and hypothalamus. In *Agpat2*^-/-^ mice, LIPIN1 levels were increased in all the brain regions. By contrast, GPAT1 in cortex and hypothalamus, AGPAT3 in hippocampus and hypothalamus, AGPAT4 in hypothalamus, and MOGAT1 in cortex, hypothalamus and cerebellum were lower in *Agpat2*^-/-^ mice. *Lepr*^db/db^ mice showed fewer and milder changes, with increased levels of GPAT1 and LIPIN1 in cerebellum, and AGPAT3 in hypothalamus.

**Conclusions and Significance:** Enzymes of glycerolipids synthesis are differentially expressed across regions of the mouse brain. Two mouse models of IR and T2D have altered gene expression of glycerolipid enzymes in the brain.

## Introduction

Type 2 diabetes (T2D) has deleterious effects on the structure and functioning of the brain [1]. These abnormalities include reduced brain and hippocampus volume and impaired attention, executive function, information processing, visuospatial construction and visual memory [1]. In mice, high-fat diet-induced insulin resistance (IR) and hyperglycemia result in loss of neurons and reduced olfactory learning [2].

Glycerophospholipid and diacylglycerol participate in the generation of second messengers, regulating apoptosis, membrane anchoring, and antioxidant capacity in the brain [3]. Preincubation of rat brain tissue sections with the radioactive phosphatidylserine (PS) increases [^3^H] α-amino-3-hydroxy-5-methyl-4-isoxazolepropionic acid (AMPA) binding to AMPA glutamate receptors without modifying [^3^H]glutamate binding to *N*-methyl-D-aspartate (NMDA) receptor, indicating that PS specifically regulate AMPA receptor binding activity [4]. Because blood brain barrier strongly limits lipid trafficking, *de novo* lipogenesis seems to the main source of glycerolipids in the brain [5].

Glycerolipids are synthesized by sequential acylation of glycerol-3-phosphate and monoacylglycerol. Glycerolipid biosynthesis starts with the formation of lysophosphatidic acid (LPA; 1-acylglycerol 3-phosphate) by glycerol-3-phosphate acyltransferases (GPATs). LPA is then acylated in *sn2* position by 1-acylglycerol-3-phosphate acyltransferases (AGPATs), forming phosphatidic acid (PA; 1,2 diacylglycerol phosphate). Phosphate group of PA is removed by phosphatidic acid phosphatases (PAPs, LIPINs), resulting in diacylglycerol (DAG) which is then acylated in *sn3* position by diacylglycerol acyltransferases (DGATs) to generate triacylglycerol (TAG) [6]. DGATs activity has been reported in very low levels in the brain [7]. Monoacylglycerol pathway starts with acylation of 2-monoacylglycerol by monoacylglycerol acyltransferases (MOGATs) to form DAG, which is further acylated by DGATs to produce TAG [7].

All the enzymes involved in glycerolipid synthesis have multiple isoforms encoded by individual genes with differential tissue expression patterns [8]. Although some of these enzymes have been reported, at the mRNA level, in whole brain homogenates [6,8], their expression across specific regions of the brain remains unknown. Furthermore, the effects of IR and T2D on the abundance of these enzymes in the brain have not been reported.

The *Agpat2* null (*Agpat2*^-/-^) mice develop generalized lipodystrophy and severe IR, diabetes, dyslipidemia and fatty liver [9] as a result of leptin deficiency [10]. Similar to *Agpat2*^-/-^ mice, mice with inactivating mutation in the leptin receptor (*Lepr*^db/db^) also develop severe IR and T2D as a result of lack of functional leptin and are obese as a result of hyperphagia and decreased energy expenditure [11,12]. In spite of the similar metabolic abnormalities between these two mouse models, we found that whereas *Lepr*^db/db^ had few and small changes in the gene expression of enzymes involved in glycerolipid biosynthesis in the brain, *Agpat2*^-/-^ mice had several changes that might be owed to specific lipid composition perturbation derived from AGPAT2 deficiency rather than to primary metabolic alterations.

## Materials and Methods

### Animals

Mice were housed 3-4 per cage, maintained on a 12 h light/12 h dark cycle and fed a standard chow diet (Prolab RMH 3000; LabDiet, St. Louis, MO) *ad libitum*. Generation of *Agpat2*^-/-^ mice has been described previously [9]. *Agpat2*^-/-^ and *Agpat2*^+/+^ mice were obtained by mating of *Agpat2*^+/-^ mice. *Lepr*^*+/+*^ and *Lepr*^*db/db*^ mice were generated by mating of *Lepr*^+/db^ mice (obtained from The Jackson Laboratory, Bar Harbor, ME). *Agpat2*^-/-^, *Agpat2*^+/+^, *Lepr*^*db/db*^ *and Lepr*^*+/+*^ adult mice (9-14 weeks) of both sexes were anesthetized by ketamine/xylazine (100:10 mg/Kg) and euthanized by exsanguination under deep anesthesia. The brain was extracted and dissected into cerebral cortex, hypothalamus, hippocampus and cerebellum and flash frozen in liquid nitrogen and stored at −80°C. All mouse procedures were reviewed and approved by the Institutional Animal Care and Use Committee (IACUC) at Pontificia Universidad Católica de Chile.

### RNA extraction and cDNA preparation

Cerebral cortex, hypothalamus, hippocampus and cerebellum were homogenized with Mixer mill from (Retsch Inc, Newtown, CT) and total RNA extracted using Trizol® reagent (Invitrogen, Carlsbad, CA) following manufacturer’s instructions. RNA was resuspended with RNAsecure solution (Invitrogen, Carlsbad, CA). RNA concentration was determined by Nanodrop spectrophotometer Nanodrop 2000 (Thermo Fisher Scientific, Waltham, MA). To evaluate the RNA integrity, 1 µg of RNA was visualized in denaturing agarose/formaldehyde gels. Residual genomic DNA was eliminated with Turbo DNA-free kit (Invitrogen, Carlsbad, CA). 2 µg of total RNA were reverse transcribed with High Capacity cDNA Reverse Transcription Kit (Applied Biosystems, Foster City, CA) following manufacturer’s instructions.

### mRNA quantification by real time PCR (RT-qPCR)

Real-time qPCR reactions were performed in Applied Biosystems StepOnePlus thermalcycler using Fast SYBR® Green Master Mix (Applied Biosystems, Foster City, CA) with 25 ng of template cDNA. Thermal cycling involved an initial step at 95°C for 20 seconds and then 40 cycles at 95°C for 3 seconds and 60°C for 30 seconds. PCR primer sequences are presented in Table 1. To determine the relative abundance of specific mRNAs across different brain regions, target gene cycle threshold (Ct) value (cut off = 30) was normalized to the Ct value of the reference gene cyclophilin (*Ppib*) using 2^-ΔCt^ method (2^-(CT target gene – CT reference gene)^). The relative mRNAs abundance of target genes in specific brains regions of wild type and *Agpat2*^-/-^ or *Lepr*^*db/db*^ mice was determined by the 2^-ΔΔCT^ method with *Ppib* as reference gene.

**Table 1:**
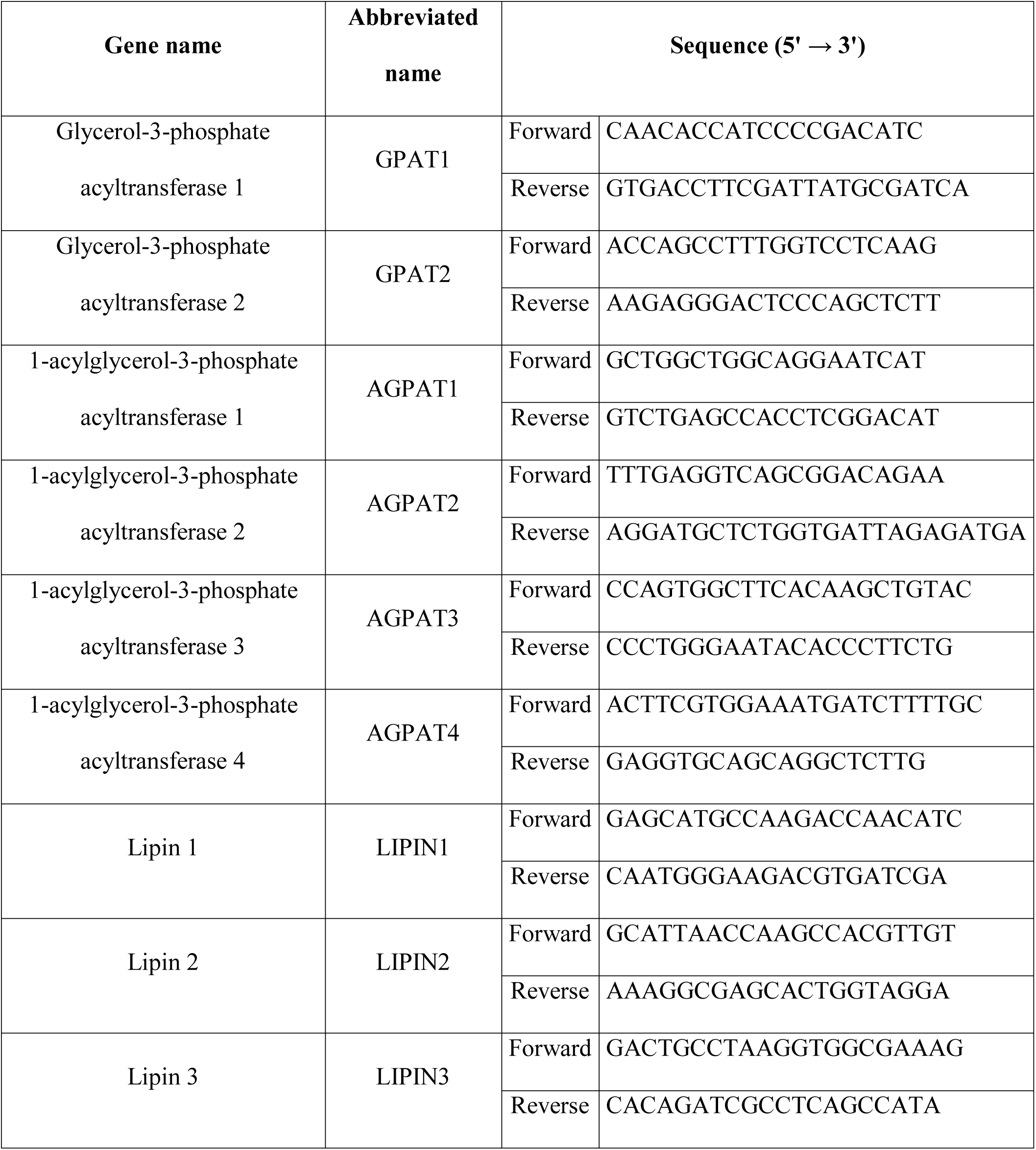
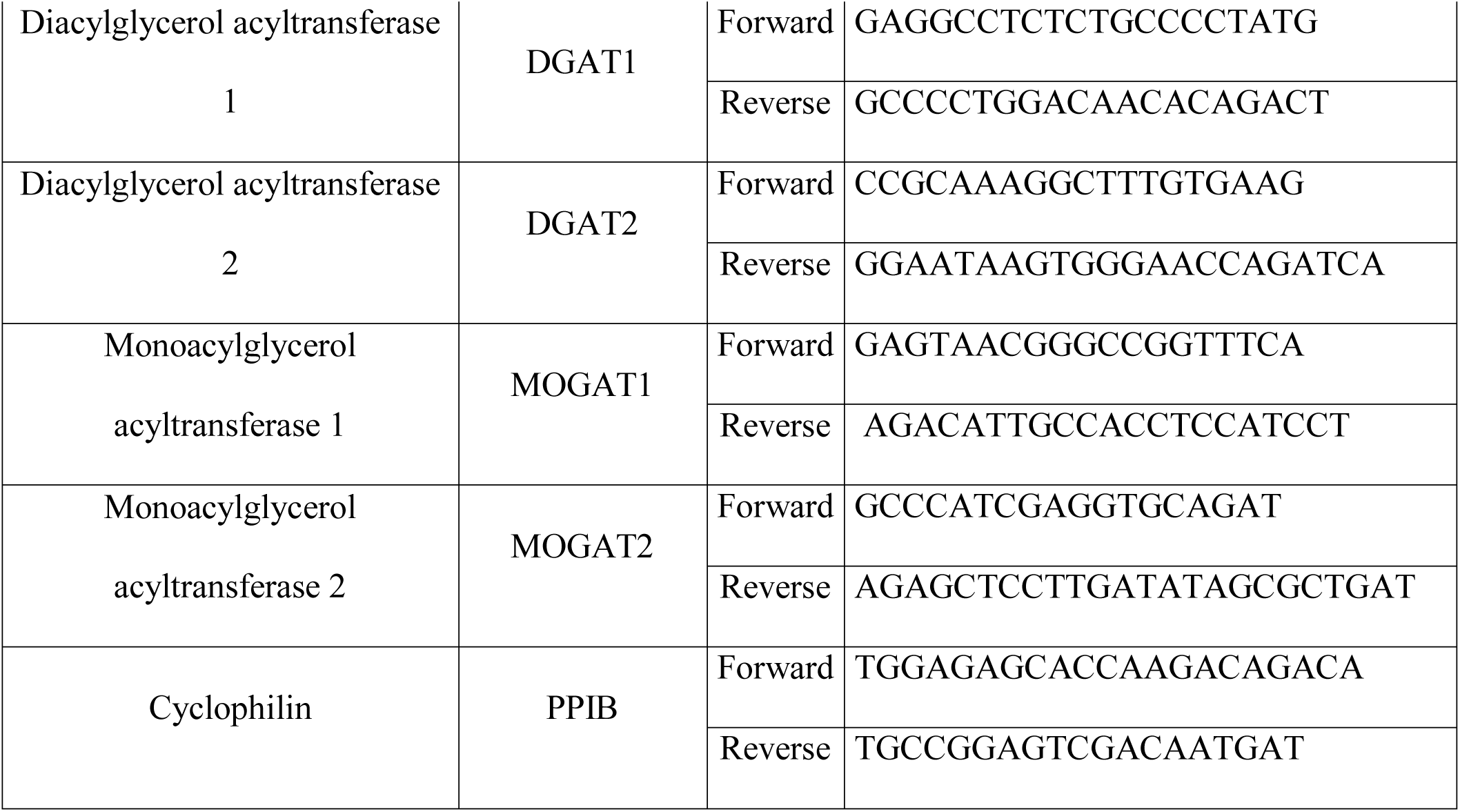
Sequence of PCR primers used for quantification of mRNAs by real time qPCR.

### Statistical analysis

For two-group comparisons, mean values were compared with two-sided Student’s t-test. P-values of <0.05 were considered significant. For experiments with more than two groups, mean values were analyzed by one-way ANOVA and Bonferroni post-hoc test for multiple comparisons. GraphPad Prism version 6.00 (GraphPad, La Jolla, CA) was used for plotting and statistical analyses.

## Results

### 1. Quantification of the mRNA levels of enzymes involved in glycerolipid synthesis in different regions of the mouse brain

A systematic gene expression analysis of the enzymes involved in glycerolipid synthesis in different regions of the brain is lacking. We studied the mRNA abundance of the main isoforms of these enzymes in the mouse cerebral cortex, hypothalamus, hippocampus and cerebellum.

GPATs mediate the first reaction in the glycerol-3-phosphate pathway, generating LPA. GPAT1 mRNA was detected in all the regions studied and its highest abundance was found in the hypothalamus, followed by the cerebellum, cortex and hippocampus (Figure 1A). GPAT2 mRNA was undetectable in all the brain regions.

**Figure 1:**
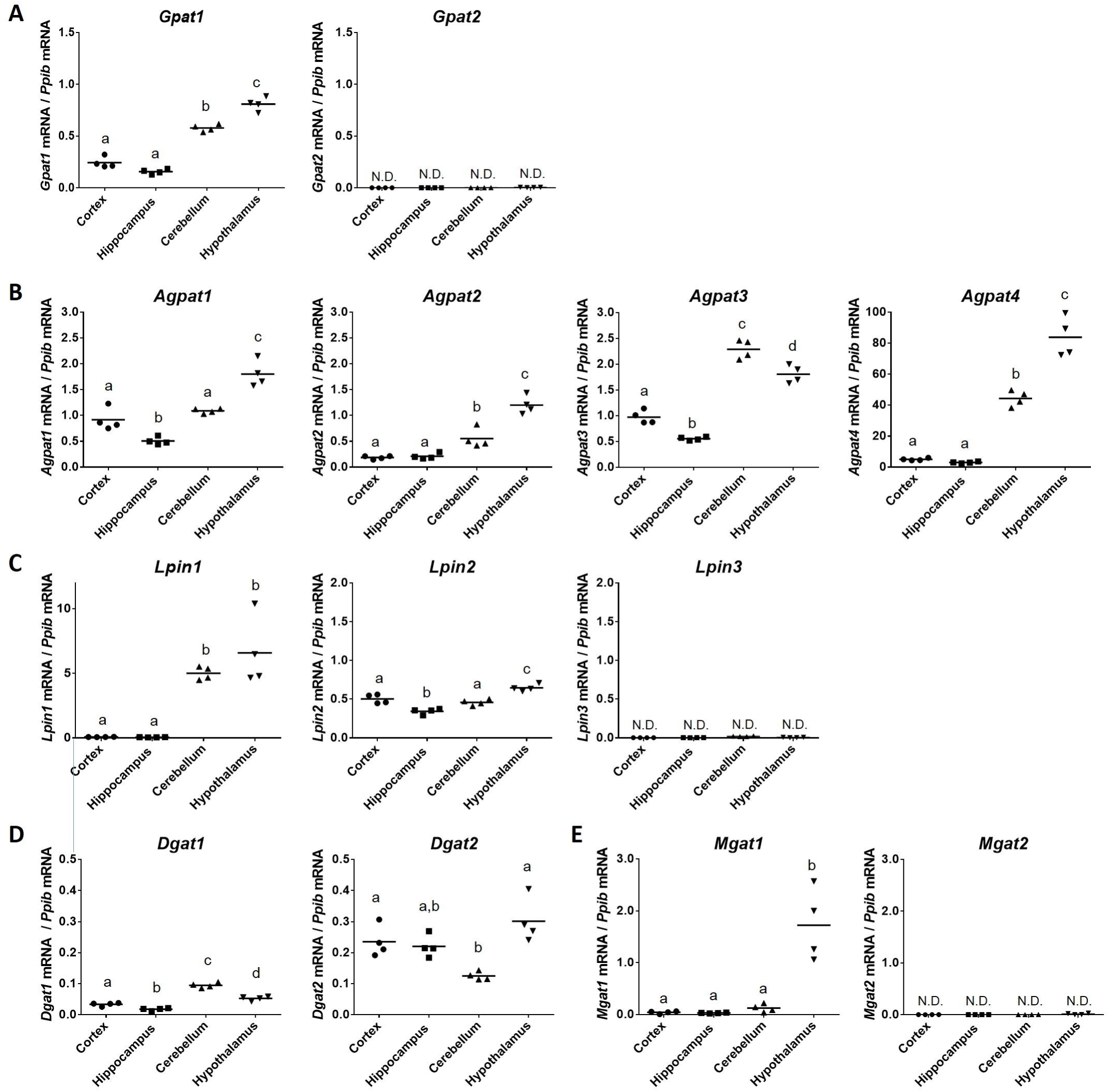
mRNA expression levels of enzymes involved in the glycerolipid synthesis in cerebral cortex, hippocampus, cerebellum and hypothalamus in wild type (C57BL/6) mice. Expression of each gene was normalized to the reference gene Cyclophilin (*Ppib*) and was calculated by the 2^-ΔCT^ method (2^-(CT target gen – CT reference gen)^). Ct value cut off was 30. Mean values were analyzed by one-way ANOVA and Bonferroni test for multiple comparisons. Statistical significance (p<0.05) is indicated by a different letter (a, b, c, d). N = 4 per group, animals were of the two sexes.

AGPATs generate PA from LPA. Among the four AGPAT isoforms that we quantified (AGPAT1-4), the most abundant was AGPAT4 whereas AGPAT2 had the lowest expression levels (Figure 1B). Interestingly, in spite of their differential abundance, AGPAT2 and AGPAT4 have the same expression pattern, with the highest expression observed in the hypothalamus, followed by the cerebellum and then cortex and hippocampus. The highest abundance of AGPAT1 was also found in the hypothalamus, but it was followed by the cortex and cerebellum, and finally the hippocampus. Unlike other isoforms, AGPAT3 has its highest expression level in the cerebellum, followed by hypothalamus, cortex and finally hippocampus.

LIPINs belong to a group of phosphatidic acid phosphatases enzymes and transform PA into DAG. The mRNA of LIPIN1 and LIPIN2 was detected in all the brain structures analyzed (Figure 1C); however, LIPIN1 mRNA abundance was significantly higher in the hypothalamus and cerebellum in comparison to cerebral cortex and hippocampus. By contrast, the highest expression level of LIPIN2 was in the hypothalamus, followed by the cortex and cerebellum, and finally the hippocampus. Interestingly, LIPIN2 was the predominant isoform in the cortex and hippocampus whereas in the cerebellum and hypothalamus the most abundant isoform was LIPIN1. LIPIN 3 mRNA was undetectable in all the studied structures.

DGAT1 and DGAT2 mRNAs were detected in all the brain regions studied, although at low levels. Of these, DGAT2 was the most abundant although its levels were similar among the hypothalamus, cortex and hippocampus but lower in the cerebellum (Figure 1D). The highest abundance of DGAT1 mRNA was in the cerebellum, followed by the hypothalamus, cortex, and hippocampus.

MOGATs produce DAG from monoacylglycerol. MOGAT1 mRNA was detected in all the studied regions of the brain, but at very low levels, with its highest abundance in the hypothalamus. MOGAT2 mRNA was undetectable (Figure 1E).

### 2. Effects of AGPAT2 deficiency and leptin receptor inactivation on the mRNA levels of glycerolipid biosynthetic enzymes in the mouse brain

To assess the effect of IR and T2D on the mRNA levels of glycerolipid biosynthetic enzymes, we studied two independent mouse models of leptin action deficiency, the lipodystrophic *Agpat2*^-/-^ and the *Lepr*^db/db^ mice.

GPAT1 mRNA levels were slightly lower in the cerebral cortex and hypothalamus of *Agpat2*^-/-^ mice in comparison with wild type (*Agpat2*^+/+^) mice but remained equivalent in all the other brain regions (Figure 2A).

**Figure 2:**
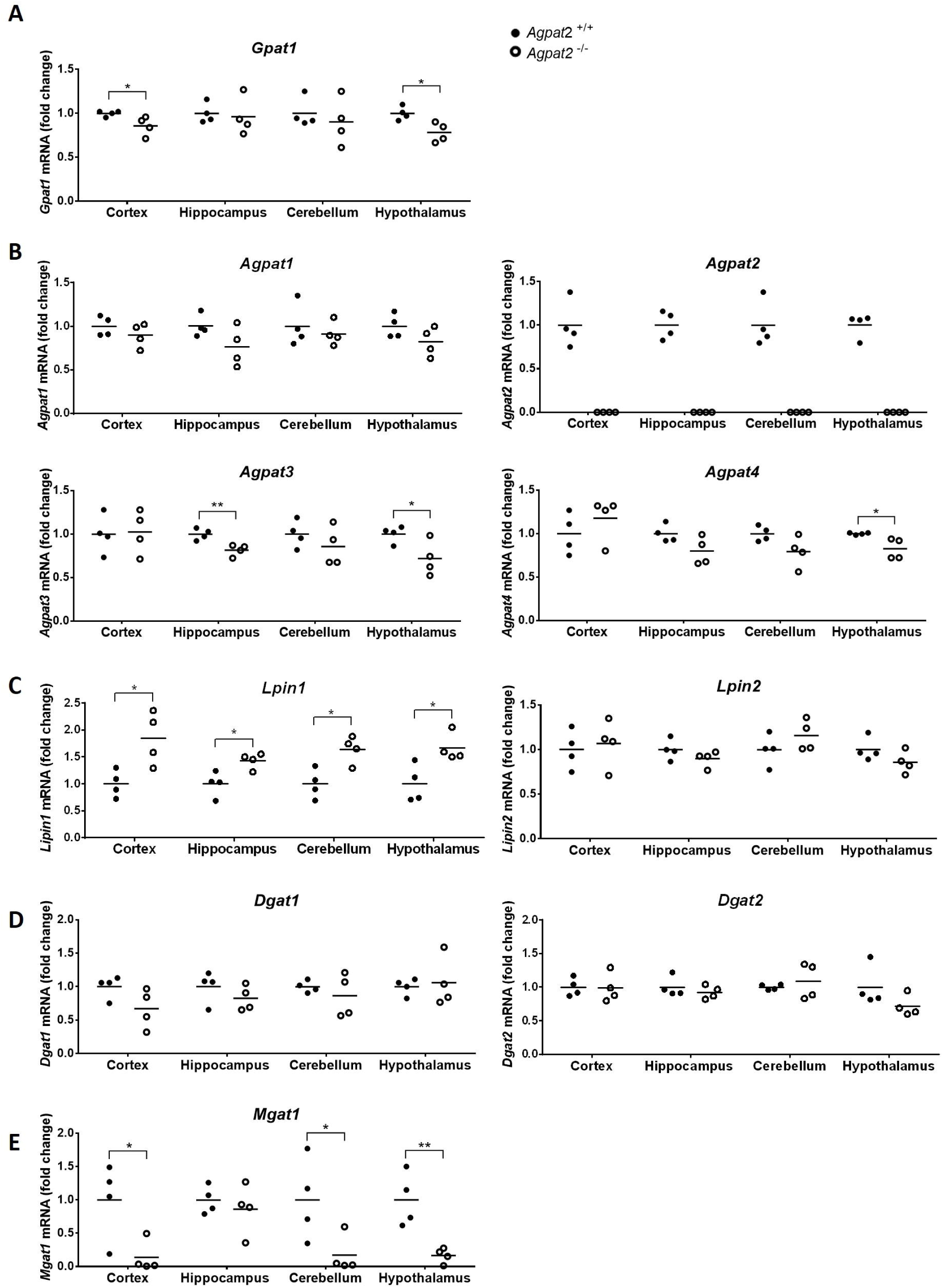
mRNA expression levels of enzymes involved in glycerolipid synthesis in cerebral cortex, hippocampus, cerebellum and hypothalamus of *Agpat2*^+/+^ and *Agpat2*^-/-^ mice. mRNA expression levels of each gene were normalized by Cyclophilin mRNA expression levels and expressed as fold-change compared to wild type (*Agpat2*^+/+^) mice, by 2^-ΔΔCT^ method. Ct value cut off was 30. * indicates statistical significance of p<0.05 and ** indicates p<0.01 by unpaired Student’s t test. N = 4 animals per group animals were of the two sexes.

No differences in AGPAT1 mRNA levels were detected in any of the studied brain regions but the abundance of AGPAT3 mRNA was lower in the hypothalamus and hippocampus of the *Agpat2*^-/-^ mice whereas AGPAT4 mRNA was only slightly decreased in the hypothalamus of these animals (Figure 2B).

LIPIN1 mRNA showed the highest differential abundance in *Agpat2*^-/-^ mice, with ∼2-fold increase in the cerebral cortex and hypothalamus and ∼1.5-fold increase in the cerebellum and hippocampus. No differences were detected in the abundance of LIPIN2 mRNA between genotypes (Figure 2C). LIPINs have reported roles in brain function. LIPIN2 deficient mice develop ataxia and decreased LIPIN1 levels as they age, resulting in altered cerebellar phospholipid composition [13]. In humans, the splicing isoform LIPIN1γ is highly expressed in the brain [14], suggesting a regulatory role on brain glycerolipid metabolism.

DGAT1 and DGAT2 mRNA levels were equivalent between *Agpat2*^-/-^ and wild type mice (Figure 2D). By contrast, MOGAT1 was strongly decreased in the cerebral cortex, hypothalamus and cerebellum of *Agpat2*^-/-^ mice but unchanged in the hippocampus (Figure 2E).

In the *Lepr*^db/db^ mice, the abundance of GPAT1 mRNA in the cerebellum was higher in comparison with wild type (*Lepr*^+/+^) mice (Figure 3A) and no differences were detected in the other regions of the brain studied.

**Figure 3:**
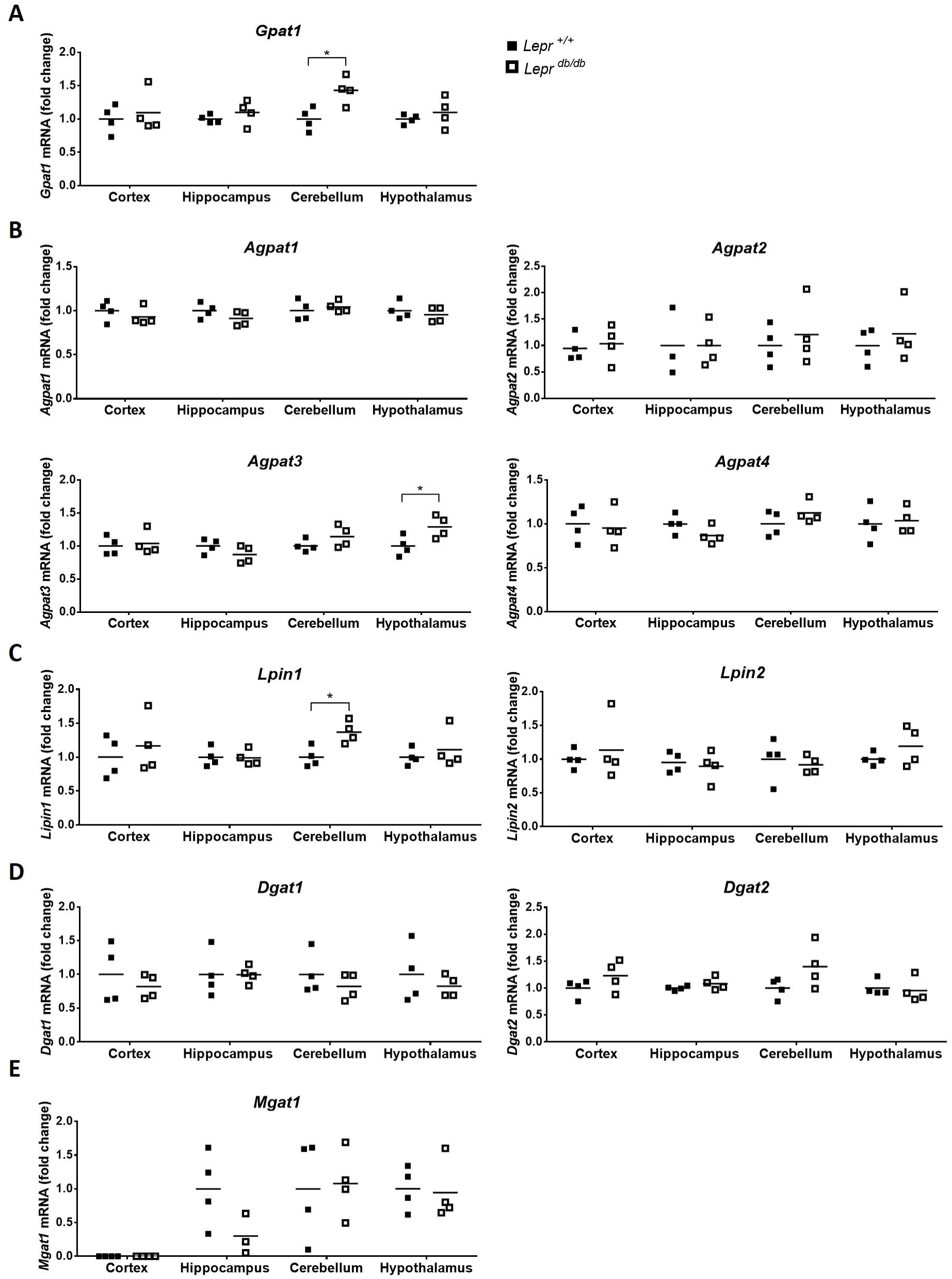
mRNA expression levels of enzymes involved in glycerolipid synthesis in cerebral cortex, hippocampus, cerebellum and hypothalamus of *Lepr*^*+/+*^ and *Lepr*^*db/db*^ mice. mRNA expression levels of each gene were normalized by Cyclophilin mRNA expression levels and expressed as fold-change compared to wild type (*Lepr*^*+/+*^) mice, by 2^-ΔΔCT^ method. Ct value cut off was 30. * indicates statistical significance of p<0.05 by unpaired Student’s t test. N = 4 animals per group, animals were of the two sexes.

Among AGPAT isoforms, only AGPAT3 mRNA levels were slightly increased in the hypothalamus of *Lepr*^db/db^ mice in comparison to wild type mice. All the other AGPATs remained unchanged (Figure 3B).

Similarly, LIPIN1 was increased only in the cerebellum of *Lepr*^db/db^ mice (Figure 3C). No differences were found in the mRNA levels of DGAT1, DGAT2 and MOGAT1 between genotypes (Figure 3D-E).

## Discussion

Here we report the mRNA levels of the key enzymes involved in glycerolipid synthesis in different regions of the mouse brain and likely impact on IR and T2D. We focused on the cerebral cortex, hypothalamus, hippocampus and cerebellum because these regions of the brain have the highest abundance of insulin receptor [15,16] and because previous studies have reported lipid composition changes in murine models of IR and T2D states in these regions of the brain [17–20].

We found that hypothalamus and cerebellum were the brain regions with the highest abundance of these enzymes (GPAT1, AGPAT1, AGPAT2, AGPAT4, LIPIN1, LIPIN2, DGAT2 and MOGAT1) at the mRNA level, suggesting important roles for the glycerolipids in the structure and/or function of these brain regions. Concordantly, it has been shown that glycerolipid metabolism is required for hypothalamic neurons controlling thermogenesis and energy expenditure in mice [21].

AGPAT4 was previously reported as the most abundant AGPAT isoform in whole mouse brain preparations [22] and its mRNA is detectable at similar levels between brain stem, cerebellum, cortex, hippocampus and olfactory bulb in mice [23]. *Agpat4*^*-/-*^ mice have impaired spatial learning and memory, in association with drastically decreased NMDA and AMPA receptor levels in the brain [24]. On the other hand, AGPAT4 is necessary for normal abundance of phosphatidylcholine, phosphatidylethanolamine and phosphatidylinositol in the brain [23]; however, whether these lipid alterations are connected to the neurological defects of the AGPAT4 deficient mice remains to be elucidated. Similarly, a recent study suggests that AGPAT1 can have roles in neuronal function, since *Agpat1*^*-/-*^ mice have abnormal hippocampal neuron development and develop audiogenic seizures [25].

The role of AGPAT2 in brain lipid composition and function has not been systematically studied. Previous studies have shown that AGPAT2 is expressed in the cerebral cortex, spinal cord, medulla, and subthalamic nucleus of the human brain [26]. However, patients with congenital generalized lipodystrophy owed to mutations in *AGPAT2* gene [27,28] and *Agpat2*^-/-^ mice [9] have no clinically evident neurological defects, suggesting that AGPAT2 is disposable for main brain functions. By contrast, neurological alterations seen in schizophrenia, depression and Alzheimer’s disease have been reported in *Lepr*^db/db^ mice [29,30]. In addition, these mice have an accelerated cognitive decline, that starts at 7 weeks of life [31–33]. Provocatively, abnormal levels of proteins involved in energy metabolism, cellular structure and neural functioning have been described in the frontal cortex and hippocampus of *Lepr*^db/db^ mice [30], indicating that abnormal metabolic regulation is correlated with abnormal neurological function in these animals. It remains to be determined whether subclinical manifestations of neurological dysfunction are present in the lipodystrophic *Agpat2*^-/-^ mice, and likely in lipodystrophic patient with mutations in *AGPAT2*, and if so whether these potential alterations are correlated with changes in glycerolipid metabolism.

Interestingly, in spite of the neurological disorders of the *Lepr*^db/db^ mice, we only found fewer and milder changes in the expression of glycerolipid enzymes: only LIPIN1 and GPAT1 were increased in the cerebellum while AGPAT3 was increased in the hypothalamus (Table 2). By contrast, in the *Agpat2*^-/-^ mice, GPAT1, AGPAT3, AGPAT4 and MOGAT1 mRNA levels were lower in the hypothalamus, whereas GPAT1 and MOGAT1 were lower in the cerebral cortex, AGPAT3 was lower in the hippocampus and MOGAT1 was lower in the cerebellum. LIPIN1 mRNA was increased in all studied brain structures in the *Agpat2*^-/-^ mice (Table 2).

**Table 2:** Summary of the changes in mRNA levels of the enzymes involved in glycerolipids synthesis in *Agpat2*^*-/-*^ and *Lepr*^*db/db*^ mice.

| Gene | *Agpat2*<sup>-/-</sup> |  | *Lepr*<sup>db/db</sup> |  |
| --- | --- | --- | --- | --- |
|  | Change | Brain regions | Change | Brain regions |
| GPAT1 | ↓ | cortex, hypothalamus | ↑ | cerebellum |
| AGPAT3 | ↓ | hippocampus, hypothalamus | ↑ | hypothalamus |
| AGPAT4 | ↓ | hypothalamus |  |  |
| LIPIN1 | ↑ | cortex, hypothalamus, hippocampus, cerebellum | ↑ | cerebellum |
| MOGAT1 | ↓ | cortex, hypothalamus, cerebellum |  |  |

The table shows only the genes that presented statistically significant changes in mRNA expression in comparison to wild type mice.
↑ indicates an increase in the mRNA expression levels with respect to wild type littermates.
↓ indicates a decrease in the mRNA expression levels with respect to wild type littermates.

The changes observed in the brain of *Agpat2*^*-/-*^ mice are in contrast with to those observed in the liver of these mice. In fact, *Agpat2*^*-/-*^ mice have increased hepatic abundance of GPAT1, AGPAT1, AGPAT3 and MOGAT1 in comparison with wild type mice [9]. In the liver, GPAT1 gene expression is potently induced by insulin by mechanisms involving sterol regulatory element-binding protein-1 (SREBP-1) [8], whereas MOGAT1 expression is promoted by PPARγ by insulin independent mechanisms [34]. The diverging regulation of GPAT1 between these two organs of the *Agpat2*^-/-^ mice can be explained by a relatively more intense IR in the brain than in the liver. Alternatively, the mechanisms regulating the gene expression of these enzymes in the brain might be different than in the liver. A recent study showed that the mRNA levels of AGPAT1–5 in liver, heart and whole brain are induced by fasting in mice [35], suggesting the physiological regulation of these enzymes is influenced by nutritional/metabolic status across different tissues. However, the physiological role of the enzymes involved in the glycerolipid biosynthesis in the brain for the metabolic adaptation to fasting remains totally unknown.

## Conclusion

In summary, we found that enzymes involved in glycerolipids synthesis are differentially expressed across regions of the brain in mice; and that two different models of IR and T2D have a differential expression pattern of some of these enzymes in different regions of the brain. *Lepr*^db/db^ had fewer and smaller changes in comparison with the *Agpat2*^-/-^ mice suggesting that the abnormal gene expression in the *Agpat2*^-/-^ mice is due to specific lipid composition perturbation derived from AGPAT2 deficiency rather than primary general metabolic alterations owed to leptin action deficiency. It remains to be studied whether these mRNA changes have an impact on the abundance of different glycerophospholipids in the brain and its role in brain functioning as a whole-body metabolic regulator.

## Acknowledgments

This work was supported by FONDECYT (Fondo Nacional de Desarrollo Científico y Tecnológico) [grant numbers 1141134 and 1181214]. The authors thank to A. Garg, A.K. Agarwal and J.D. Horton from UT Southwestern for providing the AGPAT2 deficient mice under the material transfer agreement with Pontificia Universidad Catolica de Chile and for their continual scientific advice. We also thank to Dolores Busso for providing the *Lepr*^db/db^ mice.

## Author contributions

V.C. designed and wrote the grant. L.G. designed specific aims and experiments of the study; performed the experiments and the analysis and interpretation of the data under the guidance of V.C. A.K.A. critically revised and contributed to the manuscript. The manuscript was written by L.G. and V.C.

